# *In vitro* and *in vivo* infection models reveal connection between periplasmic protease Prc and alternative peptidoglycan synthase PBP3_SAL_ in *Salmonella enterica* serovar Typhimurium

**DOI:** 10.1101/2022.12.15.520552

**Authors:** Kim Vestö, Rikki F. Frederiksen, Iina Snygg, Anna Fahlgren, Maria Fällman, Mikael Rhen

## Abstract

A hallmark in salmonellosis is the ability of the bacteria to proliferate within host cells. Most notably, *Salmonella* proliferates within professional phagocytes in a vacuolar compartment. During proliferation *Salmonella* has to build new cell wall, but how this is regulated within the intraphagosomal niche is not known. Here we show that genetically inactivating the periplasmic protease Prc, involved in cleaving peptidoglycan-processing enzymes, results in decreased fitness in macrophage-like RAW264.7 cells and in BALB/c mice, and in a decreased tolerance to redox stress. All these *prc* mutant phenotypes were conditional depending on *pbp3sal*, a recently defined paralogue for *ftsl* coding for the essential penicillin binding protein 3. These phenotypic connections between Prc and PBP3_SAL_ adds to the phenotypes governed by Prc, and possibly adds PBP3_SAL_ to the pool of target proteins involved in cell wall homeostasis that are regulated by Prc.

## Introduction

Many bacterial pathogens, such as the Gram-negative enteric bacterium *Salmonella enterica*, include an intracellular growth phase as an essential part of the infection pathogenesis (1,2). The pathogenesis of human typhoid fever is characterized by the systemic dissemination of *Salmonella* into the liver, spleen, and bone marrow after first traversing the intestine (1). Within the host the path of the bacteria likely includes at least two intracellular growth phases; transcytosis of the bacteria through the intestinal epithelial cell layer (3,4) and replication in phagocytes (5). As the causative agents of typhoid and paratyphoid fever, respectively *S*. Typhi and *S*. Paratyphi, are strictly human adapted whereby studies on typhoid and paratyphoid fever have relied on systematic murine salmonellosis models to provide detail to their pathogenesis (6). Accompanying studies chiefly relying on *S*. Typhimurium have clearly corroborated intracellular replication in phagocytes and a use of dendritic cells as transport vehicles for systemic spread (5,7–9). Due to the ability of phagocytes to produce reactive oxygen species, *Salmonella* possesses several modes of adaptation to oxidative stress as a virulence trait to ensure intracellular replication (10,11).

When proliferating within macrophages *Salmonella* synthesizes new peptidoglycan, as indicated by peptidoglycan synthesis genes being transcribed in the intramacrophagal niche and that intracellular salmonellae can be treated with peptidoglycan synthesis inhibitors (13). An integral part of cell wall synthesis in proliferating bacteria is to create the septum, i.e. the structure that will divide the bacterium into two daughter cells. In *Enterobacteriaceae*, the peptidoglycan synthase responsible for the completion of the septum synthesis is the essential penicillin-binding protein 3 (PBP3) (14,15). However, recently Castanheira *et. al*. (16) discovered that *Salmonella* possesses a non-essential homologue to PBP3, named PBP3_SAL_. Furthermore, Castanheira *et. al*. (16) showed that an environment mimicking the intracellular compartment *Salmonella* replicates in enhances production of PBP3_SAL_. In contrast, expression of the canonic PBP3 was not induced in PBP3_SAL_-inducing *in vitro* medium or in murine RAW264.7 cells (17). This suggests a possible redundancy in the function of PBP3 that has evolved for *Salmonella* to be able to adapt its peptidoglycan synthesis while remaining intracellular (16).

An additional enzyme connected to peptidoglucan turnover is the periplasmic protease Prc (also known as Tsp), entwined in regulation of cell wall metabolism due to its ability to proteolytically process the C-termini of PBP3 (18–20), and to degrade both murein endopeptidase MepS (also known as Spr) (21) and lytic the transglycosylase MltG in *Escherichia coli* (22). Yet, at least in *E. coli*, a non-enzymatic outer membrane lipoprotein, NlpI, acts as a cofactor for Prc (21). Additionally *prc* was identified in a screen to be part of a locus involved in survival of *Salmonella* within primary peritoneal murine macrophages (23). This led us to further detail the role of Prc in the fitness of *Salmonella* during infection, and for adaptation to host innate defense effectors.

## Material and Methods

### Bacterial strains

The bacterial strains used in this study were *S*. Typhimurium line SR-11. The Δ*pbp3sal*-mutation and cloning to create the pBAD30::*prc* plasmid were constructed as described previously (24). All strains used in the study and the associated primers to create mutants are in Table 1 and Table 2.

**Table 1.**
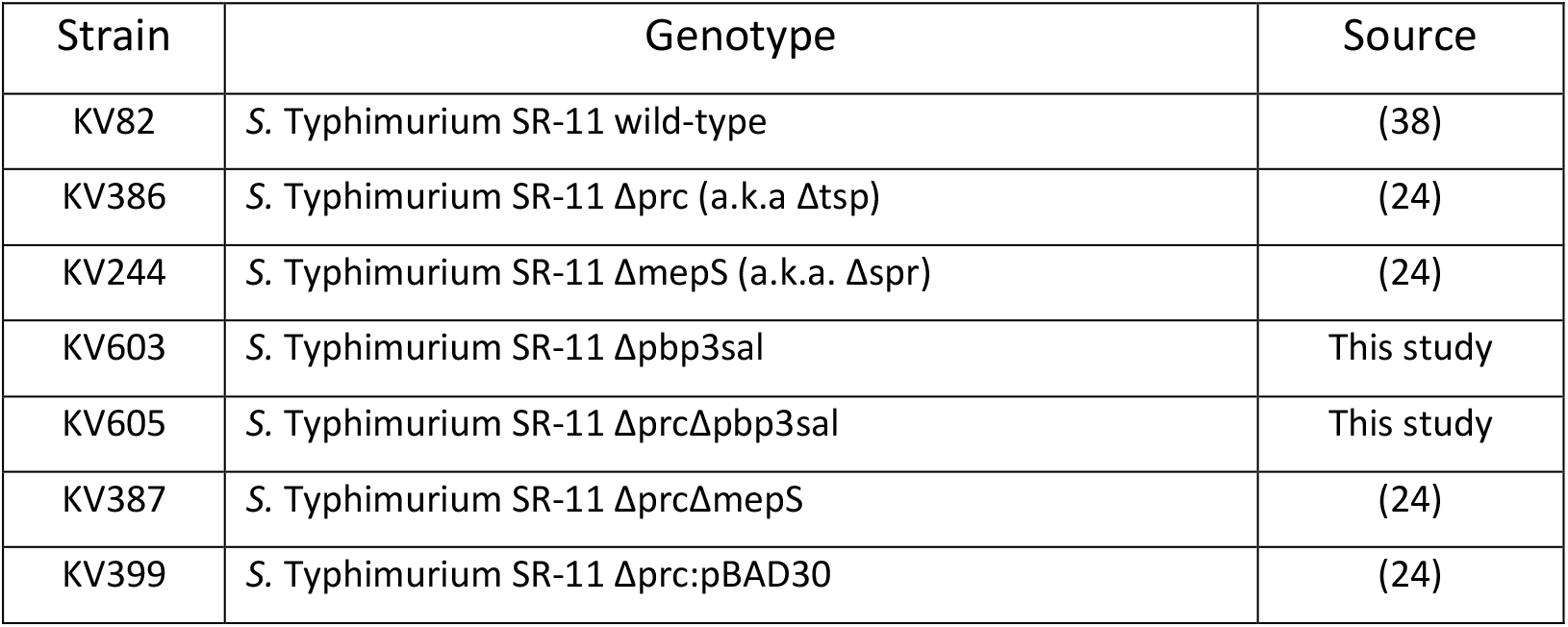
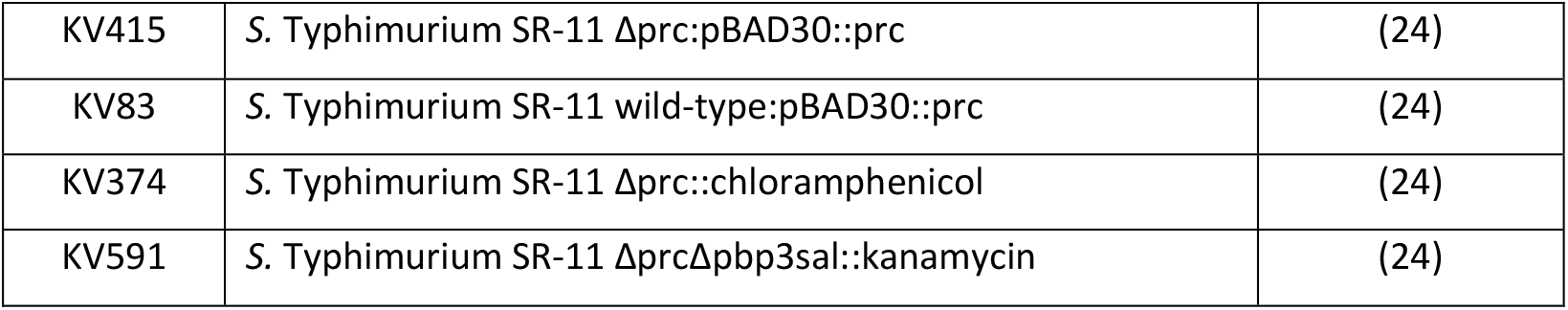
Strains used in study.

**Table 2.**
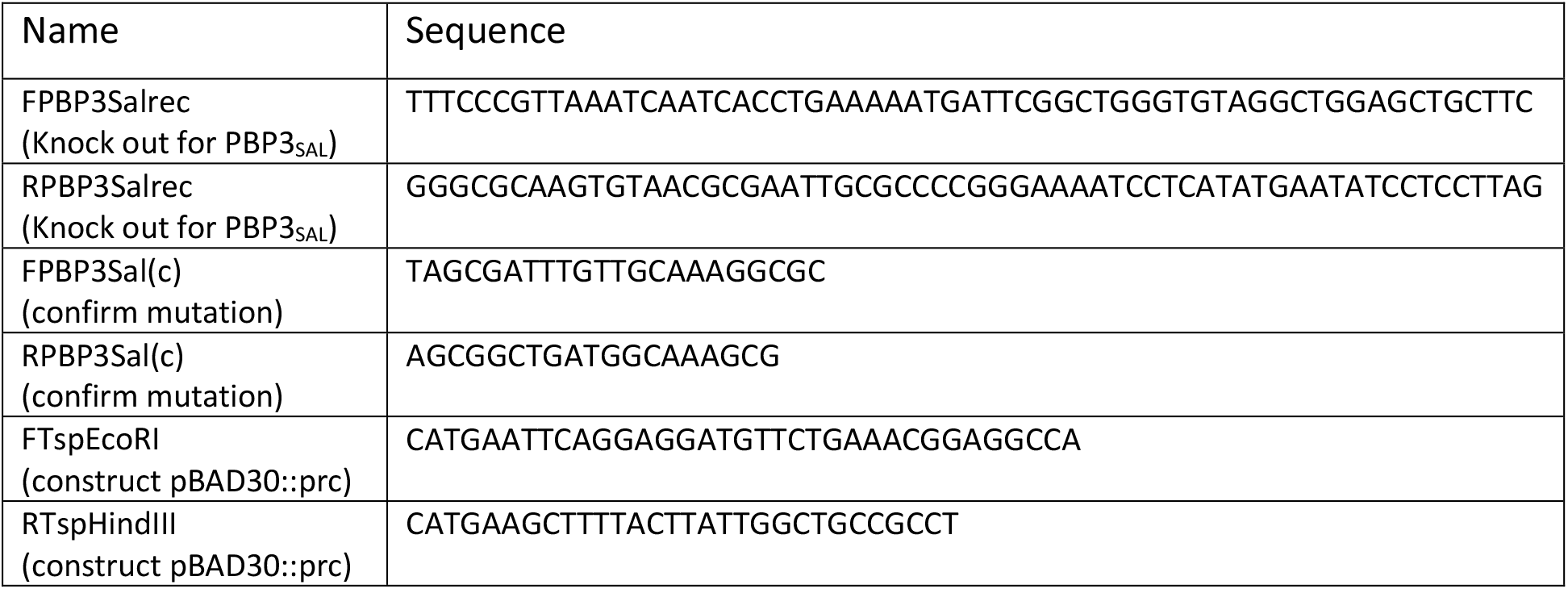
Primers used in study.

### Cell culture

The murine macrophage cell line RAW264.7 (25) was purchased from American Type Culture Collection ATCC TIB-71) and cultured at 37°C under 5% CO_2_ in maintenance media consisting of RPMI 1640 (Gibco, 21875-034) supplemented with 10% heat-inactivated FBS (Fetal Bovine Serum, Gibco), 2mM L-glutamine (Gibco, 25030-024), 20mM HEPES (Sigma-Aldrich, H0887), and 10μg/ml gentamicin (Sigma-Aldrich, G1272).

### Gentamicin protection assay

For the gentamicin protection assay the RAW264.7 cells were seeded into 24 well plates (Sarstedt) in maintenance medium the day before infection at approximately 10^5^ cells per well.

For infection, *Salmonella* were first cultured overnight in 2ml LB (10mg/ml NaCl, Sigma-Aldrich) in 15ml Falcon tubes at 220rpm and 37°C. Next day the cultures were diluted 1:100 in fresh 2ml of LB in 15ml Falcon tubes and cultured for 2 hours at 220rpm and 37°C. The OD_600nm_ of the cultures was determined using Ultrospec 1000 (Pharmacia Biotech) and the volume of culture needed to be added to 10ml of infection media to have a final concentration of 10^6^ colony-forming units (CFUs)/ml was calculated using the formula ((0.484/OD_600nm_) x 21) where the output is the amount of μl needed for the 10ml. For genetic complementation experiments, LB was supplemented with 100μg/ml ampicillin (Sigma-Aldrich) and 0.02% L-arabinose (Sigma-Aldrich) in each incubation.

Bacteria were added to 10ml of infection media (RPMI 1640 and 20mM HEPES) in three duplicates at a MOI of 10 after removing the maintenance media. One set of duplicate wells was used to enumerate the CFUs taken up by the RAW264.7 cells two hours post-infection. The second set of duplicate wells was used to enumerate the CFUs for overnight growth of the bacteria within the RAW264.7 cells 20 hours post-infection. A third set of duplicate wells was identical to the second but with addition of 10ng/ml interferon-γ (IFN-γ; Sigma-Aldrich) to the media.

Following addition of bacteria to the wells, the plate was centrifuged at 500 rpm (48 relative centrifugal force, Eppendorf Centrifuge 5810R) with slow start and stop for 10 minutes in order to aid with internalization of the bacteria. This was followed by one hour incubation at 37°C under 5% CO_2_. After the incubation the infection media was replaced by maintenance media supplemented with 50μg/ml gentamicin, and incubated for 1 hour at 37°C under 5% CO_2_ in order to kill bacteria not internalized.

One set of duplicate wells were washed twice with 1ml PBS followed by incubation at room temperature with 1ml of 0.5% sodium deoxycholate (Sigma-Aldrich) in PBS for 5 minutes in order to lyse macrophages and release the bacteria. After 5 minutes the wells were vigorously pipetted and 100μl of the content was transferred into Eppendorf tubes containing 900μl of PBS in order to perform serial dilution and plating on LB agar plates to enumerate the CFUs of internalized bacteria with the CFU values denoted “uptake”. The second set of duplicate wells the killing media was replaced by maintenance media. The third set was replaced with maintenance media supplemented with 10μg/ml IFN-γ and incubated for 20 hours at 37°C under 5% CO_2_. Next day the second and third sets were treated as above to release the bacteria and the CFUs enumerated from the LB agar plates denoted as “overnight growth”.

The growth of intracellular bacteria was done by taking the ratio of CFU of overnight growth divided by the uptake for each mutant or wild-type to get a value for the proliferation within the macrophages. This value was then normalized to the wild-type value in order to reveal possible fitness defects in intracellular proliferation in comparison to wild-type and denoted “normalized fitness”.

### Drop-on-lawn tests

Sensitization phenotypes were conducted essentially as described by Vestö *et al*. (24) however replacing vancomycin with either hydrogen peroxide or ascorbic acid (both purchased from Sigma Aldrich).

### Competition infection in mice

For the *in vivo* infection model a competition experiment in 8 week old female BALB/c mice (Charles River) was performed. Bacteria were grown overnight at 37°C at 100rpm in LB (5mg/ml NaCl, VWR) with 50μg/ml kanamycin or 50μg/ml chloramphenicol (Sigma-Aldrich) supplemented for the mutants. The following day, overnight cultures were diluted 1:20 in LB and grown for 4 hours to approximately and OD_600nm_ of 1.0 in Erlenmeyer flasks. The cultures were diluted to 10^4^ CFUs/ml in PBS, mixed in a 1:1 ratio between wild-type and mutant, and 100μl of this mixture was administered by intraperitoneal injection into mice. To confirm the inoculum, CFUs were enumerated following serial dilution and plating on LB agar plates followed by incubation overnight at 37°C.

Three days post infection the mice were anesthetized and sacrificed by cervical dislocation and liver, spleen, and gallbladder recovered and homogenized (GentleMACS dissociator) in cold PBS. The homogenates were serially diluted in cold PBS and plated on LB agar plates followed by incubation overnight at 37°C. The following day plates with 100-250 colonies were replica plated onto LB agar plates and LB agar plates supplemented with either kanamycin or chloramphenicol, depending on the mutant that was competed against wild-type, in order to determine the proportion of wild-type to mutant in the population. The replica plating was done by placing a sterile velvet pad on the plates and transferring the velvet to fresh plates as noted above. The calculation of the competitive index was performed by calculating the ratio between the CFUs of the wild-type and the CFUs of the mutant when recovering them from the organs after normalizing the wild-type and the mutant to their respective input CFUs.

For the experiment the mice were housed in accordance with the Swedish National Board for Laboratory Animals guidelines. All animal procedures were approved by the Animal Research Ethics Committee of Umeå (Dnr A 27-2017). Mice were allowed to acclimate to the new environment for one week before experimental onset.

### Statistics

One-way ANOVA with Dunnett’s correction was performed for assays comparing multiple mutants with wild-type at the same time. Student’s t-test was done when an assay involved only two different bacteria i.e. a mutant and a wild-type. The statistics were performed using GraphPad Prism v6.0g (GraphPad Software, Inc., USA).

## Results

### Prc is needed for full fitness during overnight growth in RAW264.7 cells

To study the role of Prc in *Salmonella* during infection we generated a *prc* deletion mutant in line SR-11 lacking *prc* and performed *in vitro* and *in vivo* infection assays. When studying phenotypes of the Δ*prc*-mutant in our *in vitro* infection model using murine macrophage RAW264.7 cells, we observed that the fitness of the Δ*prc*-mutant was significantly lower than that for the wild-type following an overnight incubation (Fig. 1A). This difference between the fitness of the Δ*prc*-mutant compared to wild-type was even more pronounced when the RAW264.7 cells were treated with interferon-γ (IFN-γ) prior to the overnight incubation (Fig. 1A). The gene *prc* is likely part of an operon (26). Hence, to confirm that the fitness defect of the Δ*prc*-mutant was due to the lack of Prc we complemented the mutant by expressing *prc in trans* resulting in a partly, yet significant, complementing the fitness defect (Fig. 1B).

**Figure 1.**
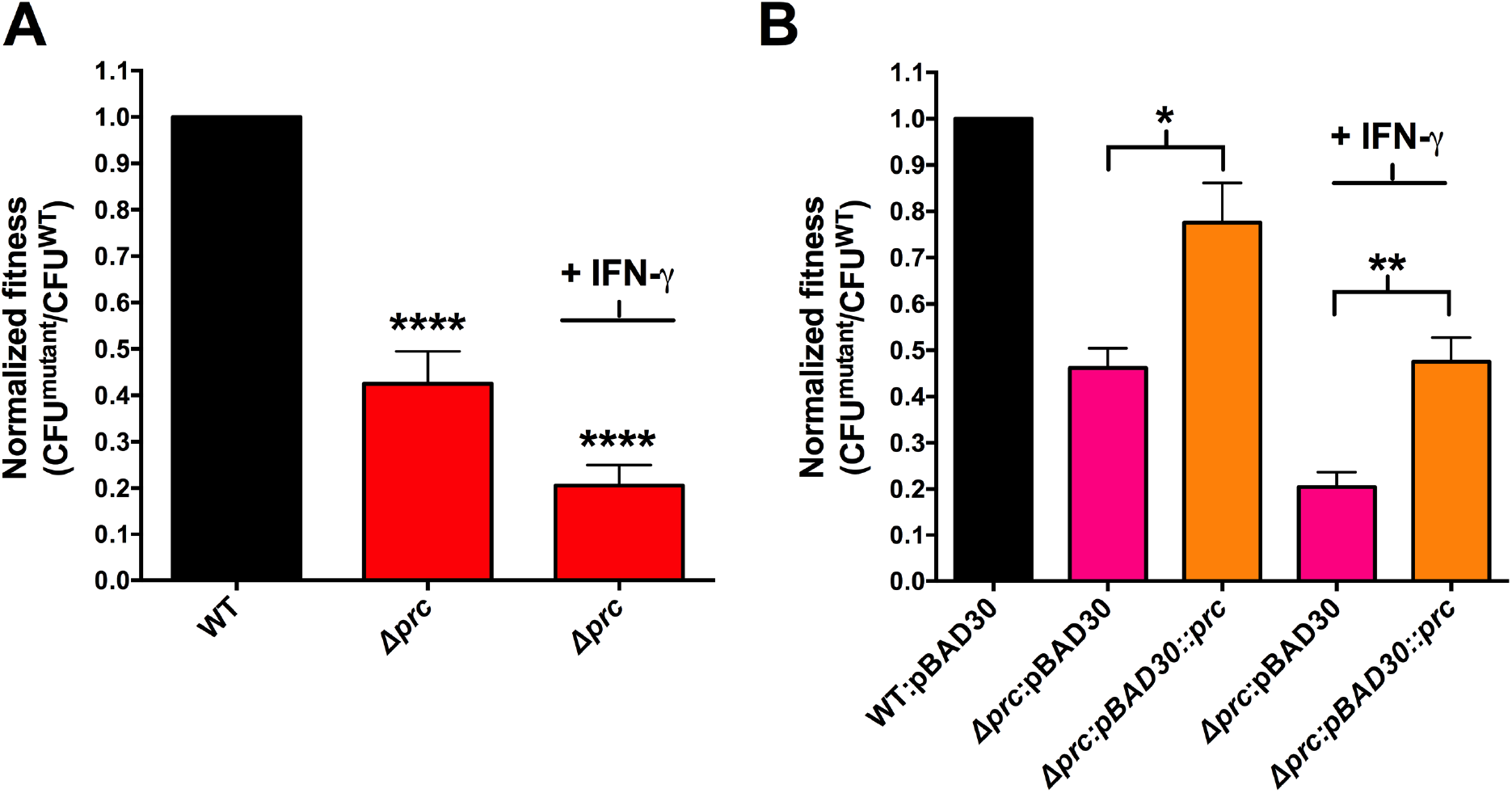
*In* vitro infection assay of murine macrophage RAW264.7 cells using the gentamicin protection assay. Each of the mutant constructs are normalized towards its respective wild-type, meaning that the non-IFN-γ treated samples are normalized towards a non-IFN-γ-treated wild-type and IFN-γ-treated mutants against IFN-γ-treated wild-type. The wild-type is represented as a single column due to the wild-type being normalized against itself in both non-treated and IFN-γ-treated condition. Each experiment was repeated at least three times. A) Normalized fitness of the Δ*prc*-mutant in comparison to wild-type (WT) in infection of RAW264.7 cells as defined by the ratio of CFU yield following overnight growth as compared to the CFU of bacteria taken up by the RAW264.7 cells normalized to WT in both non-activated and IFN-γ-activated RAW264.7 cells. B) Normalized fitness of Δ*prc*-mutant complemented using pBAD30::*prc* plasmid in both non-activated and IFN-γ-activated RAW264.7 cells. IFN-γ also affected wild-type bacteria with the wild-type in non-IFN-γ-activated RAW264.7 cells growing to a fitness ratio on average of 72 (fitness ratio = CFU_overnight growth_/CFU_uptake_), while in the IFN-γ-activated RAW264.7 cells the fitness ratio is 12 i.e. the IFN-γ-activation of the RAW264.7 cells inhibits the overnight growth of the wild-type 6-fold. One-way ANOVA with Dunnett’s correction was performed with statistical significance denoted as * = p <0.05, ** = p <0.01, **** = p<0.0001.

### The effect on fitness by Prc in RAW264.7 cells is PBP3_SAL_-dependent

Phenotypes associated with genetically knocking out the periplasmic protease Prc are thought to be due to the accumulation of the murein endopeptidase MepS as lack of *prc* results in a 10-fold increase in the amount of MepS in the bacteria (21). Additionally, phenotypes of bacteria lacking *prc* can be suppressed by further removal of *mepS*, a mechanism that has been shown for various phenotypes in both *E. coli* and *Salmonella* (21,24,27,28). To reveal if altered MepS levels contributed to the fitness phenotype in RAW264.7 cells of the Δ*prc*-mutant we generated a double mutant lacking both these genes. When the resulting Δ*prc*Δ*mepS*-mutant was subjected to infection of RAW264.7 cells, the fitness defect still remained (Fig. 2). This lack of suppression was not due to the Δ*mepS*-mutation itself being detrimental for the growth of *Salmonella* in the RAW264.7 cells (Fig. 2).

**Figure 2.**
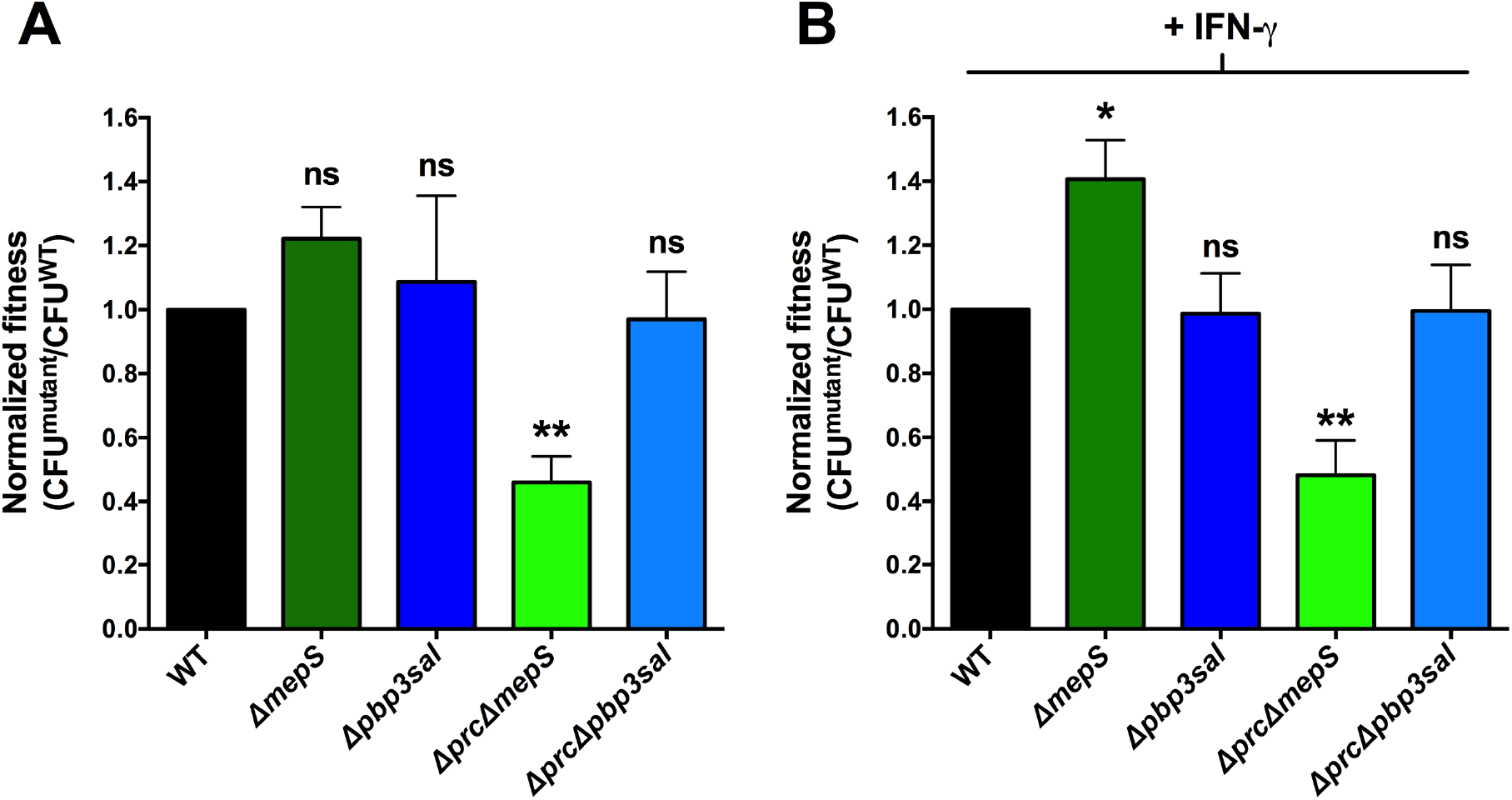
*In* vitro infection assay of murine macrophage RAW264.7 cells using gentamicin protection assay. Normalized fitness to wild-type (WT) of various mutants in both non-activated (A) and IFN-γ-activated (B) RAW264.7 cells following overnight growth. Each experiment repeated at least three times. The bar for wild-type is normalized against itself and the mutants are normalized against the wild-type exposed to the same condition i.e. mutants in non-activated RAW264.7 cells are normalized against wild-type grown in non-activated RAW264.7 cells and likewise IFN-γ-activated normalized against IFN-γ-activated wild-type. As stated in figure 1 the IFN-γ-activation of the RAW264.7 cells inhibits the overnight growth of the wild-type 6-fold meaning that the wild-type in (A) has a 6-fold yield to the wild-type in (B). One-way ANOVA with Dennett’s correction was performed with statistical significance denoted as ns = not significant, * = p <0.05, ** = p <0.01.

As the removal of *mepS* failed to suppress the fitness defect of the Δ*prc*-mutant in the RAW264.7 cells we opted for a candidate approach in trying to find a suppressor mutation for the Δ*prc*-mutant fitness defect. Another protein that has been shown to functionally interact with Prc, albeit in *E. coli*, is the peptidoglycan synthase PBP3. Prc has been shown to cleave PBP3 (18–20), yet further studies on the phenotypic and genetic associations of Prc and PBP3 are cumbersome due to the essentiality of PBP3 (29). However, due to the recent characterization of a paralogue to PBP3, named PBP3_SAL_, by Castanheira *et. al*. (16) we postulated that PBP3_SAL_ activity could possibly be another direct or indirect target of Prc. When we removed *pbp3sal* from the Δ*prc*-mutant we observed that the fitness defect was completely suppressed during overnight growth in both non-IFN-γ-activated and IFN-γ-activated RAW264.7 cells, an effect not due to the lack of *pbp3sal* itself resulting in higher proliferation than wild-type (Fig. 2).

### Prc is needed for tolerance to oxidative and reducing compounds in a PBP3_SAL_-dependent manner

Replication of *S*. Typhimurium in phagocytic cells is associated with exposure to oxidative stress (10,11). To explain the decreased fitness of the Δ*prc*-mutant in the phagocytic RAW264.7 cell line, we tested the mutant for sensitization to oxidative stress. In this indeed the mutant appeared less tolerant to hydrogen peroxide *in vitro*, as compared to the wild-type (Fig. 3A).

**Figure 3.**
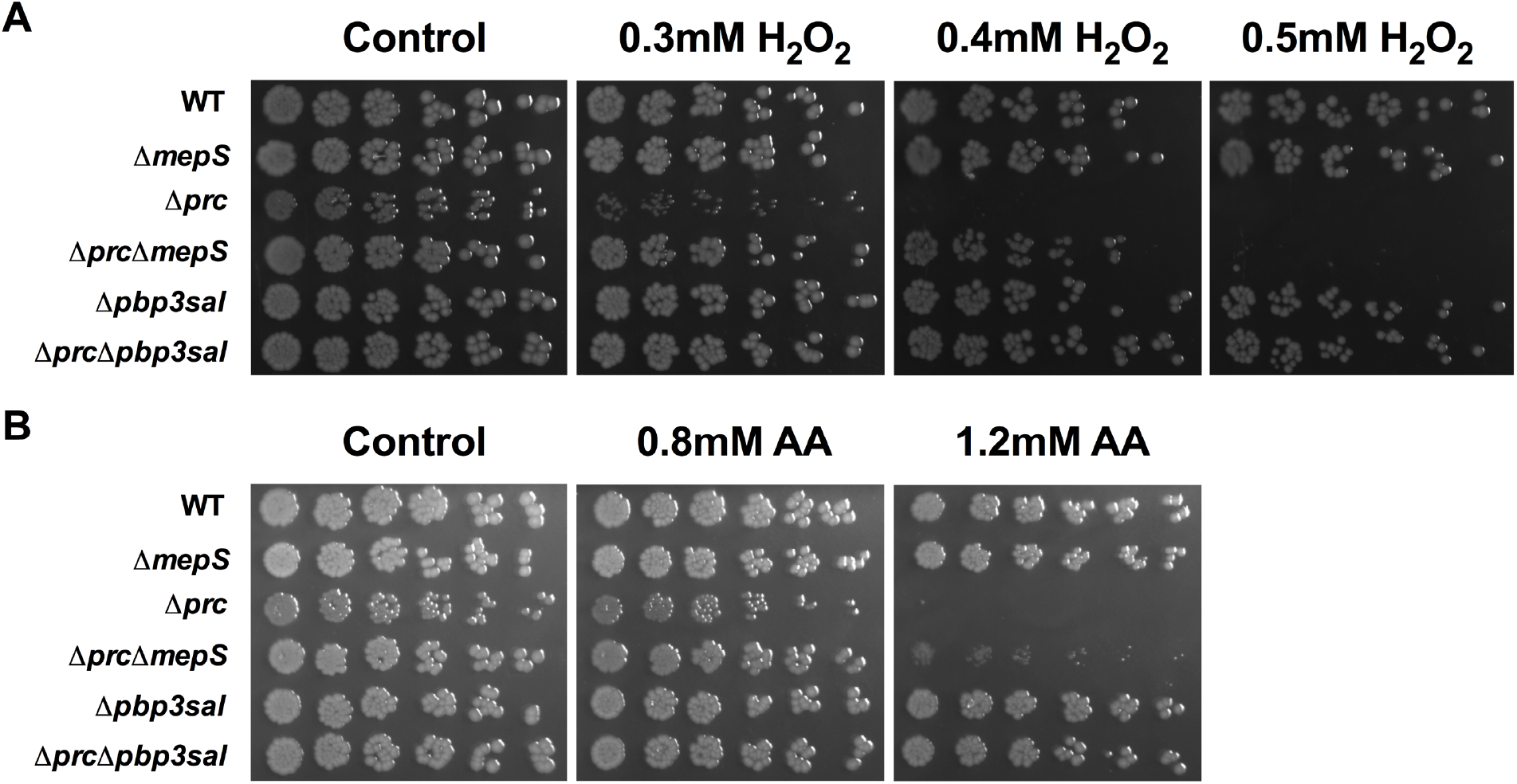
Drop-on-lawn test probing redox sensitization. Bacterial suspensions (10^5^ CFU per ml) were diluted 1:2 to 1:64 in a succession and 5μl drops were placed on the nutrient plates from the right to the left. Plates were supplemented with either hydrogen peroxide or ascorbic acid (AA) with non-supplemented plates and the wild-type (WT) serving as controls.

We have previously shown that RAW264.7 cells infected with S. Typhimurium is heterogeneous with a proportion of hypoxic cells containing large amounts of bacteria, and with another normoxic proportion of cells containing few or no bacteria (30). Thus, we tested whether the Δ*prc*-mutant was sensitized to reducing stress. Indeed, the mutant was sensitized to ascorbic acid (Fig. 3B).

Significantly, sensitizations to either hydrogen peroxide or ascorbic acid were fully suppressed by deleting *pbp3sal*, but only partially suppressed by a *mepS* deletion (Fig. 3A & 3B). Thus, the decrease in growth yield of the mutant could relate to its inability to survive in the hypoxic cell population, otherwise permissive for replication of the wild-type.

### Prc is needed for full fitness in BALB/c mice in a PBP3_SAL_-dependent matter

To extend our findings from cell cultures and in vitro redox stress measurements, we performed a competition infection in BALB/c mice. In this we observed that the Δ*prc*-mutant had a substantially decreased fitness as it was readily outcompeted by the wild-type when enumerating CFUs from liver, spleen and gallbladder, as evident by the Δ*prc*/WT competition resulting in a competitive index of <1 for all the organs (Fig. 4). Similarly to the *in vitro* model of RAW264.7 cells the removal of *pbp3sal* from the Δ*prc*-mutant also reverted the fitness defect of the Δ*prc*-mutant in the BALB/c mice as seen by the fact that the Δ*prc*Δ*pbp3sal*-mutant yielded equal CFUs from liver, spleen and gallbladder as the wild-type (Fig. 4).

**Figure 4.**
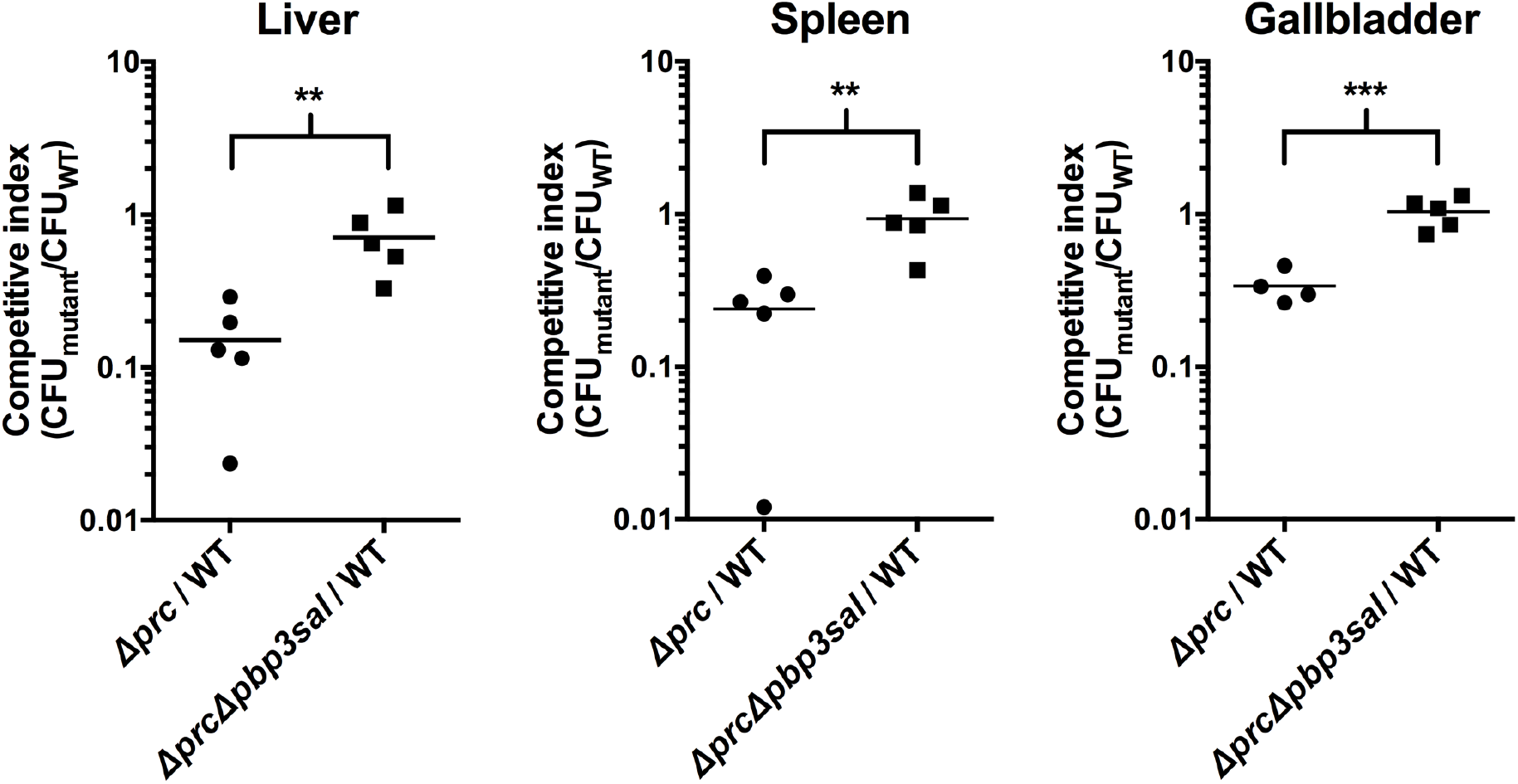
Competitive infection in BALB/c mice. Mutant and wild-type (WT) bacteria were mixed in 1:1 ratio and inoculated intraperitoneally into 8-week-old female BALB/c mice. Each data point indicates an individual mouse. At day 3 of the infection the ratio of mutant to wild-type were enumerated from liver, spleen and gallbladder in order to define the competitive index for each mutant in relation to the wild-type. A competitive index of <1 means that the recovered CFU from the organs was higher for the wild-type than for the mutant and a competitive index of >1 means that there was a higher CFU recovered for the mutant than for the wild-type following infection. Student’s t-test was performed with statistical significance denoted as ** = p<0.01 and *** = p<0.001.

## Discussion

In the past Fields *et. al*. (9) created a set of transposon mutants in *S*. Typhimurium unable to grow in macrophages from which Bäumler *et. al*. (23) subsequently described the loci needed for in intramacrophagal survival. At the time seven out of 30 mutations were affecting known genes, one of which was one coding for periplasmic protease Prc (23). Here we confirmed this observation by further extending it into an *in vivo* model while also genetically confirming via complementation the need for *prc* in an *in vitro* cell culture model. We furthermore define new redox-associated phenotypes for a *prc* mutant that could explain attenuation of a *S*. Typhimurium *prc* mutant in infection models.

While Prc was initially described as a phage tail specific periplasmic protease in *E. coli*, Prc has other substrates (31), such as MepS coding for a muramyl endopeptidase (21). The concomitant regulation of the MepS levels by Prc is of great importance as the phenotypes associated with a Δ*prc*-mutation, mainly in *E. coli*, are seemingly due to accumulation in MepS levels (21,32). Indeed, examples of published Δ*prc*-mutant phenotypes in *E. coli*, including growth defect at an elevated temperature (27) and hypo-osmotic media (21), as well as decreased sensitivity to mecillinam (28) have been shown to be suppressed by inactivation of *mepS*. Furthermore, the novobiocin sensitization of a *prc* mutant was recently shown by us to be suppressed by a *mepS* mutant (24). Deleting *mepS*, in *S*. Typhimurium did however not suppress the virulence-associated attenuations of the *prc* mutant, with an exception of a partial complementation of the fitness loss under redox stress. However, the attenuations in virulence traits were suppressed by deleting *pbp3*sal coding for an alternative PBP3 in *Salmonella*, and with orthologues in *Citrobacter* spp. and *Enterobacter spp*. (16). Thus, at leats in S. Typhimurium, may possess MepS independent acitivties.

Relevant for the peptidoglycan turnover, Prc also degrades the muramyl endopeptidase MltG and PBP3 (22). Thus the cause of the observed virulence-associated attenuations of the *S*. Typhimurium Δ*prc* mutant could be multifold. However that one can restore the pathogenicity-associated fitness costs by deleting *pbp3sal* points to a functional, direct or indirect, interaction between Prc and PBP3_SAL_. In *E. coli* PBP3 is reported to form both homo- and PBP3/PBP1B heterodimers (33). Considering the rather homologous central protein domains in PBP3 and PBP3_SAL_, possibly also PBP3_SAL_ could undergo oligomerization. In *S*. Typhimurium line SR-11 used here and in line SL1344 used by Casteinheina *et al*. (16), expression of *pbp3sa*l is increased under phagocyte-like growth conditions (34). Thereby, in the absence of Prc, concomitant accumulation of periplasmic proteins involved in cell wall synthesis could cause abbreviations in functional protein complexes, possibly involving PBP3_sal_, eventually resulting in decreased fitness.

Regardless, our data strongly implicate a role of an inherited intact peptidoglycan dynamics for virulence fitness characters in *S*. Typhimurium. This would be in concert with several recent publications. For example, treatment of human typhoid fever, caused by *S*. Typhi with the β-lactam ceftriaxone to some extent associates with relapses (13). Interestingly, in this respect Castenheira *et al*. (17) demonstrated that PBP3_SAL_ is needed for establishment of relapses after ceftriaxon treatment in a murine infection model. As for other pathogens, PBPs have been implicated in virulence of *Brucella melitensis* (35) and in redox tolerance of *Listeria monocytogenes* (36), while exposing *Staphylococcus aureus* to sublethal vancomycin affects virulence gene expression (37).

